# CX3CR1 Engagement by Respiratory Syncytial Virus Leads to Induction of Nucleolin and Dysregulation of Cilia-related Genes

**DOI:** 10.1101/2020.07.29.227967

**Authors:** Christopher S. Anderson, Tatiana Chirkova, Christopher G. Slaunwhite, Xing Qiu, Edward E. Walsh, Larry J. Anderson, Thomas J. Mariani

## Abstract

Respiratory syncytial virus (RSV) contains a conserved CX3C motif on the ectodomain of the G-protein. The motif has been indicated as facilitating attachment of the virus to the host initiating infection via the human CX3CR1 receptor. The natural CX3CR1 ligand, CX3CL1, has been shown to induce signaling pathways resulting in transcriptional changes in the host cells. We hypothesize that binding of RSV to CX3CR1 via CX3C leads to transcriptional changes in host epithelial cells. Using transcriptomic analysis, the effect of CX3CR1 engagement by RSV was investigated. Normal human bronchial epithelial (NHBE) cells were infected with RSV virus containing either wildtype G-protein, or a mutant virus containing a CX4C mutation in the G-protein. RNA sequencing was performed on mock and 4-days-post-infected cultures. NHBE cultures were also treated with purified recombinant wild-type A2 G-protein. Here we report that RSV infection resulted in significant changes in the levels 766 transcripts. Many nuclear associated proteins were upregulated in the WT group, including Nucleolin. Alternatively, cilia-associated genes, including CC2D2A and CFAP221 (PCDP1), were downregulated. The addition of recombinant G-protein to the culture lead to the suppression of cilia-related genes while also inducing Nucleolin. Mutation of the CX3C motif (CX4C) reversed these effects on transcription decreasing nucleolin induction and lessening the suppression of cilia-related transcripts in culture. Furthermore, immunohistochemical staining demonstrated decreases in in ciliated cells and altered morphology. Therefore, it appears that engagement of CX3CR1 leads to induction of genes necessary for RSV entry as well as dysregulation of genes associated with cilia function.

**Importance:** Respiratory Syncytial Virus (RSV) has an enormous impact on infants and the elderly including increased fatality rates and potential for causing lifelong lung problems. Humans become infected with RSV through the inhalation of viral particles exhaled from an infected individual. These virus particles contain specific proteins that the virus uses to attach to human ciliated lung epithelial cells, initiating infection. Two viral proteins, G-protein and F-protein, have been shown to bind to human CX3CR1and Nucleolin, respectively. Here we show that the G-protein induces Nucleolin and suppresses gene transcripts specific to ciliated cells. Furthermore, we show that mutation of the CX3C-motif on the G-protein, CX4C, reverses these transcriptional changes.

## Introduction

Respiratory Syncytial Virus causes respiratory disease in humans(1). Initial infections occur usually within the first year of life and continually throughout childhood and adulthood(2, 3), (4, 5). RSV infection initiates in the upper airways and can be found in the lower airways during severe disease(6). Symptoms are generally mild, manifesting in stages similar to the common cold (e.g. rhinorrhea, cough, sneezing, fever, wheezing) but can manifest as viral pneumonia during severe disease which occurs mostly in young children and elderly adults (7). The economic impact of RSV in the United States is estimated at over a half a billion dollars and the World Health Organization has listed RSV as a public health priority(8, 9).

RSV contains a negative-sense, single-stranded, genome. The RSV genome encodes 11 separate proteins using host translation machinery(1). Two proteins, G and F, facilitate attachment and viral penetration into the host cell(10). The RSV G-protein contains a CX3C motif that has been implicated as an attachment motif of RSV(11–15). The CX3C motif is similar to that found in fractalkine (CX3CL1) and has been shown to bind the host fractalkine receptor, CX3CR1, although perhaps in a manner unique from fractalkine(16). RSV F-protein has been implicated in both attachment and entry into the host epithelial cell. The attachment motif of F is still unknown, but attachment to the host protein via Nucleolin, has been well reported(17).

RSV primarily targets host epithelial cells in the lungs(18), although many cell types are infectible(19–21). Multiple host proteins have been indicated in RSV attachment including CX3CR1, Nucleolin, ICAM-1, Heparan sulfate, chondroitin sulfate(22). The physiological location of some these proteins on the apical lung epithelium has not been completely explored, although CX3CR1 have been shown to exist on apical surface of human lung tissue(12, 15). CX3CR1 has been shown as a co-receptor for other viruses, including HIV(23), and variations in the CX3CR1 allele has been shown to increase susceptibility to HIV-1 infection and progression to AIDS(24). Although the exact mechanism behind attachment and fusion of RSV to the host is still being investigated, host CX3CR1 appears to play a role in RSV infection.

CX3CR1 engagement by its ligand CX3CL1 (fractalkine) has been shown to regulate cellular transcription via its heterotrimeric Gαi protein. Specifically, the binding of CX3CR1 by CX3CL1 has been shown to increase numerous signaling molecules including several different secondary messengers and transcription factors(25). If attachment of RSV to CX3CR1 results in host epithelial cell transcriptional changes is unknown. Here we use primary human differentiated epithelial cell cultures to evaluate the effect of CX3CR1 engagement by RSV.

## Methods

### Virus Propagation

Four RSV strains were used for the NHBE infection: A2, A2CX4C mutant with the an alanine A^186^ insertion in the CX3C motif (^182^CWAIAC^187^) of G protein, recombinant rA2 line 19F (r19F), and r19FCX4C mutant (provided by Dr. M.L. Moore, formerly at Emory University and presently with Meissa Vaccines, Inc., South San Francisco, CA). The prototype A2 strain of RSV (ATCC, Manassas VA) was propagated in HEp-2 cells after serial plaque purification to reduce defective-interfering particles. A stock virus designated as RSV/A2 Lot# 092215 SSM containing approximately 5.0 × 10^8^ PFU/mL in sucrose-stabilizing media and stored at −80 °C for inoculation. This stock of virus has been characterized and validated for upper and lower respiratory tract replication in the cotton rat model. The A2CX4C, r19F and r19FCX4C viruses were grown in HEp-2 cells at 0.1 or 0.01 multiplicity of infection (MOI) to reduce defective-interfering particles, purified through a sucrose cushion, and stored at −80 °C. For inoculation, the virus was thawed, diluted to a final concentration with PBS (pH 7.4 w/o Ca2+ or Mg2+), kept on ice, and used within an hour.

### NHBE Cells

Normal human bronchial epithelial cells healthy adult patients were kindly provided by Dr. C.U. Cotton (Case Western Reserve University), expanded, and cultured as described (26). Briefly, cells were plated on a layer of mitomycin C (Sigma) treated 3T3 mouse fibroblast feeder cells and grown until 70% confluency in the 3:1 mixture of Ham’s F12/DMEM media (HyClone) supplemented with 5% FBS (Sigma), 24 μg/mL adenine (Sigma), 0.4 μg/mL hydrocortisone (Sigma), 5 μg/mL Insulin (Sigma), 10 ng/mL EGF (Sigma), 8.4 ng/mL Cholera toxin, and 10 μM ROCK1 inhibitor Y-2763 (Selleck Chemical LLC). After expansion, cells (passage 3 or 4) were plated on Costar Transwell inserts (Corning Inc., Corning, NY), grown until confluency, then transferred to air-liquid interface where cells were maintained in 1:1 mixture of Ham’s F12/DMEM media supplemented with 2% Ultroser G (Pall Biosepra, SA, Cergy-Sainte-Christophe, France) for 4 weeks until they were differentiated. Differentiation was confirmed by transepithelial resistance measurements >500 ohms, and flow cytometry staining for FoxJ1 (eBioscience) and acetylated α-tubulin (Life Technologies).

### NHBE Inoculation

The differentiated NHBE cultures were inoculated with RSV MOI of 0.1 or 0.3 as determined by infectivity titration in HEp-2 cells. The NHBE were washed with PBS and virus in PBS or PBS alone was added to the apical surface of the cells and incubated for 1 h at 37 °C, the virus inoculum aspirated, the apical surface washed with PBS and fresh media added to the basolateral compartment. The RSV-infected NHBE were incubated for 4 days at 37 °C and 5% CO_2,_ and then cell lysates were collected by washing cells twice with PBS and adding 100ul of RNA lysis buffer (Qiagen) to each well.

### RNA Isolation

RNA was extracted, DNAse treated, and purified using a Qiagen RNeasy kit (QIAGEN). The RNA was reverse transcribed into cDNA using an iScript cDNA synthesis kit (Bio-Rad) following the manufacturer’s instruction.

### Real-Time PCR

Semi-quantitative PCR was carried out on a 7500 Fast Real-time PCR system (Applied Biosystems) using Power SYBR Green PCR master mix (Applied Biosystems). The CT values were normalized using control PPIA CT values from the same samples. Top genes were validated using qPCR performed on samples from a separate experiment. Primer sequences: NCL Primer FWD GAACCGACTACGGCTTTCAAT, NCL Primer REV AGCAAAAACATCGCTGATACCA; CFAP221 Primer FWD GGGTTGTTCGCAATCAAGAAGA, CFAP221 Primer REV GTTGAAACCGTTTTTGTGCCC; CC2D2A Primer FWD CAAGCAGCGAGGTCCAAAG, CC2D2A Primer REV GGCTCTGTGCCAAATTCAGTC.

### RNA Sequencing

Purified RNA was sequenced using the HiSeq 2500. Gene with low reads counts were filtered as previously described(27). Filtered reads were aligned to the GRCh38 reference genome. The RSV A2 virus genome (GenBank: KT992094.1) was used for virus transcript alignment. Gene counts were normalized and log transformed (R 3.4.4, rLog package). Multiple comparisons were corrected using the Benjamini-Hochberg method (R 3.4.4, “p.adjust()” command).

### Statistical Models

Significant association between transcript levels and condition, both marginal association (mock infection vs RSV infection) and viral-transcript-level adjusted models (wildtype and CX4C mutant), was determined by linear regression (R 3.4.4. *lm* function). The wildtype and CX4C mutant comparisons were adjusted in the statistical model by using the eigenvalue for each sample derived using principal component analysis (FractalMineR) of 11 RSV gene transcript levels. Resulting p-values were adjusted for multiple testing using Benjamini-Hochberg method (R 3.4.4. *p.adjust* function). P values less than 0.05 was considered significant.

### Pathway Analysis

Gene symbols with unadjusted p-values of less than 0.05 was used for pathway analysis. Pathway analysis of gene sets were performed using ToppGene functional analysis software(28) and genes were referenced against the Gene Ontology (GO) Biological Process GO Term.

## Results

### Transcriptional Changes after RSV Infection

The transcriptional profile of normal human bronchial epithelial cells after infection with RSV was determined. Infection by RSV both increased and decreased gene transcripts relative to mock infected (Table 1, Supplemental Document 1). RSV infection significantly increased 558 gene transcripts. Infection resulted in the induction of transcripts encoding antiviral enzymes (OAS1, OAS2, CMPK2, HERC5, HERC6), antiviral peptides (RSAD2), interferon response genes (RIG-I, STAT1, STAT2, MX1, IFIT1, IFIT2, IFI44L, IRF1, OASL), mucin genes (MUC5AC, MUC5B, MUC15, MUC21, MUC13), and stat inhibitor genes (SOCS1, SOCS53), chemokines/cytokines (CX3CL1, CCL5, CXCL9, CXCL8, CXCL10, TGFA), interleukins and their receptors (IL1A, IL15, IL15R IL22RA1, IL2RG, IL18R1), Major Histocompatibility Complex genes (HLA-A, HLA-B, HLA-C, HLA-E, HLA-F, HLA-DMA, HLA-DRB1), Type-II helper T cell associated genes (PSENEN), toll-like receptors (TLR2), and innate immune cell differentiation/survival genes (BATF2, JAGN1). Both pro-apoptotic (CASP1, CASP4, CASP7, CASP8) and anti-apoptotic (Birc3, BCL2L13, BCL2L14, BCL2L15) transcripts were increased during infection. The 2′-O-ribose cap methyltransferase (CMTR1), a protein required for viral RNA cap snatching and shown to decrease interferon type I, was also increased after infection. Taken together, these results demonstrate many genes were upregulated in bronchial epithelial cells in response to RSV.

**Table 1.**
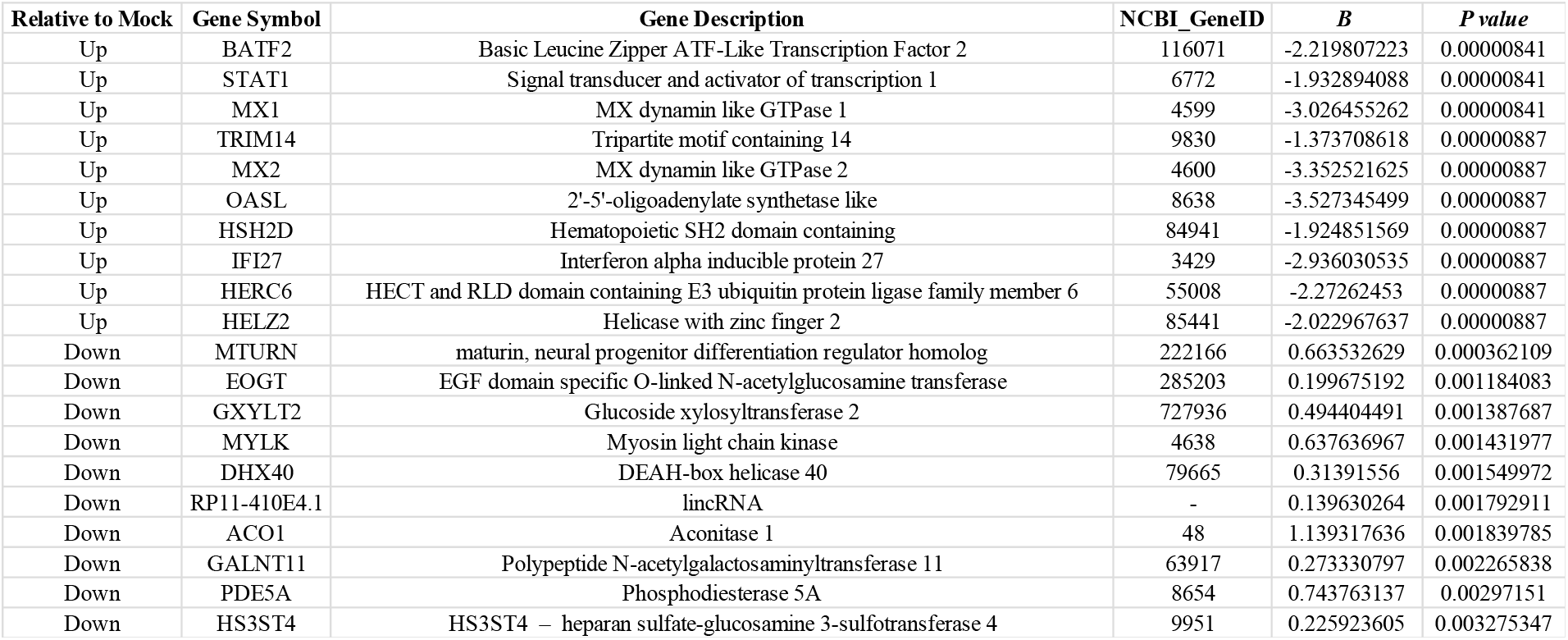
Mock vs Wildtype RSV Infection Differentially Expressed Genes

Pathway analysis of significantly upregulated genes (Mock vs RSV) demonstrated 1033 significantly enriched biological processes (Figure 1A, Supplemental Document 2). The most significantly enriched biological processes included defense processes, responses to foreign objects, innate immune responses, and cytokine-associated responses. Innate immune responses were also found to be enriched. Taken together, we found that cellular defense and immune system biological processes were associated with increased viral transcript after RSV infection.

**Figure 1.**
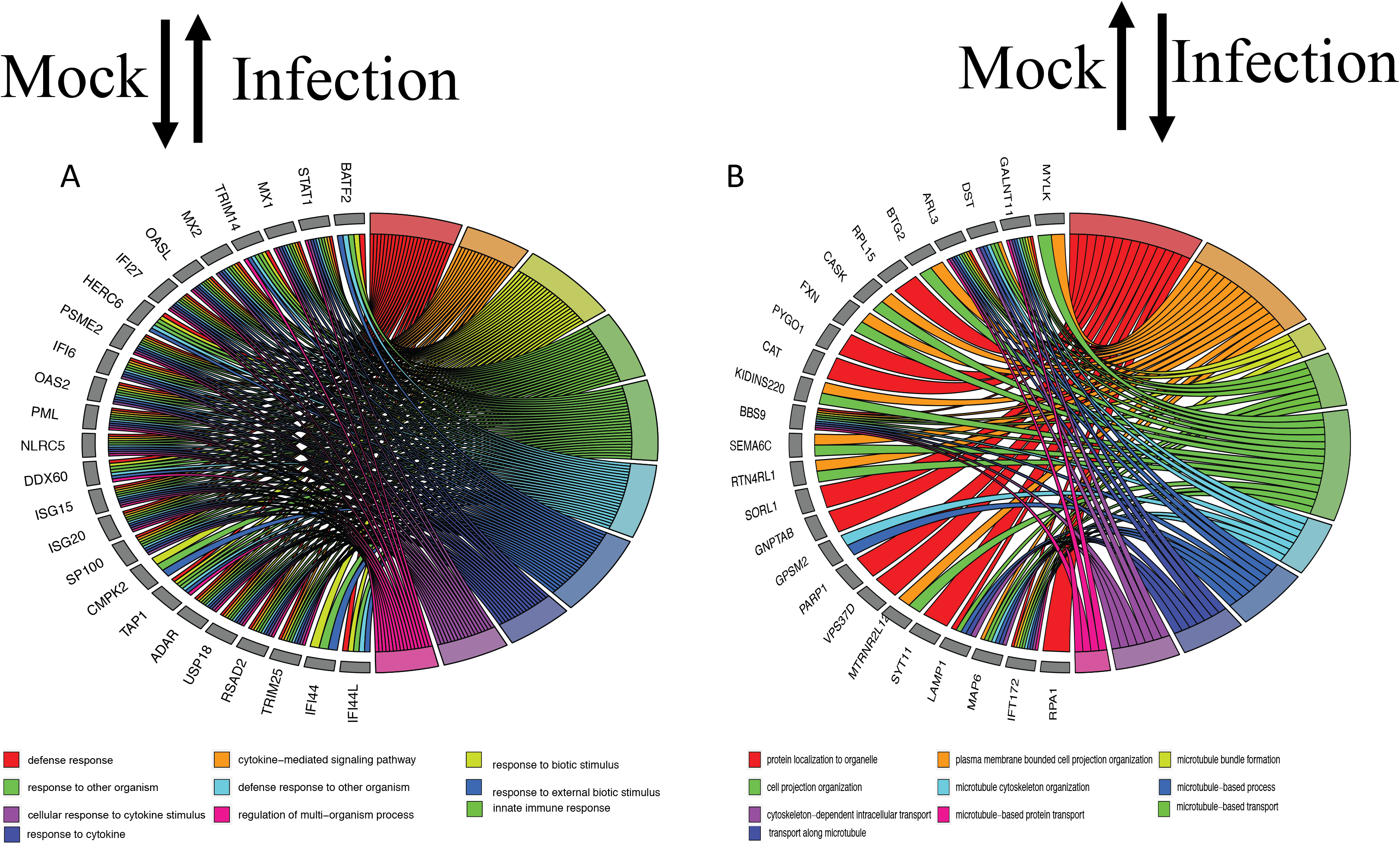
Changes in Biological Processes After RSV Infected. (A) Top ten biological processes, and associated genes, significantly enriched in transcripts increased after RSV infection. (B) Top ten biological processes, and associated genes, significantly enriched in transcript decreased after RSV infection. Colors represent the biological processes distinguished in the legend.

RSV infection also resulted in the reduction of 208 transcripts relative to mock infection with many transcripts associated with cell morphology (Table 1, Supplemental Document 1). This included microtubule-associated protein transcript levels (MAP6, MAPT, MAST1) and myosin transcript levels (MYO6, MYO9A, MYO3A, MY05A) were decreased after infection. We found many cilia associated genes (CC2D2A, PCDP1), lysosomal genes (LAMP1), leukotriene B4 hydrolase (LTA4H), WNT signaling transcripts (MCC, WNT2B), and RNA helicases (DHX40) to also be decreased during infection. Interestingly, the heparan sulfate sulfotransferase (HS3ST4) was among one of the most significantly associated down-regulated transcripts.

Pathway analysis of differentially expressed down-regulated genes identified 110 enriched biological processes (Figure 1B, Supplemental Document 3). Down-regulated genes were associated with biological process related to cellular morphology, including cell projection. Microtubule and cellular transplant related biological processes were also enriched for down-regulated genes including microtubule bundle formation, cytoskeletal transport, and protein localization.

### CX3C Specific Transcriptional Changes

Infection with an A2 RSV virus containing a mutation in the CX3C motif (CX4C) resulted in a number of significant transcriptional changes relative to the wildtype virus (Table 2, Supplemental document 2). Five host transcripts were significantly decreased after mutant virus infection compared to wild-type virus infections. NCL (Nucleolin), the eukaryotic nucleolar phosphoprotein, was the most significantly decreased gene in the mutant virus infection compared to wildtype. The adenylate kinase, AK2, an apoptotic activator; the Golgi homeostasis protein, TMEM199, recently identified to be essential in influenza A virus infection; and the transcriptional coactivator (SNW1) which can bind the vitamin D receptor binding domain and retinoid receptors to enhance many gene expression pathways were also decreased after infection with the mutated compared to the wildtype virus.

**Table 2.**
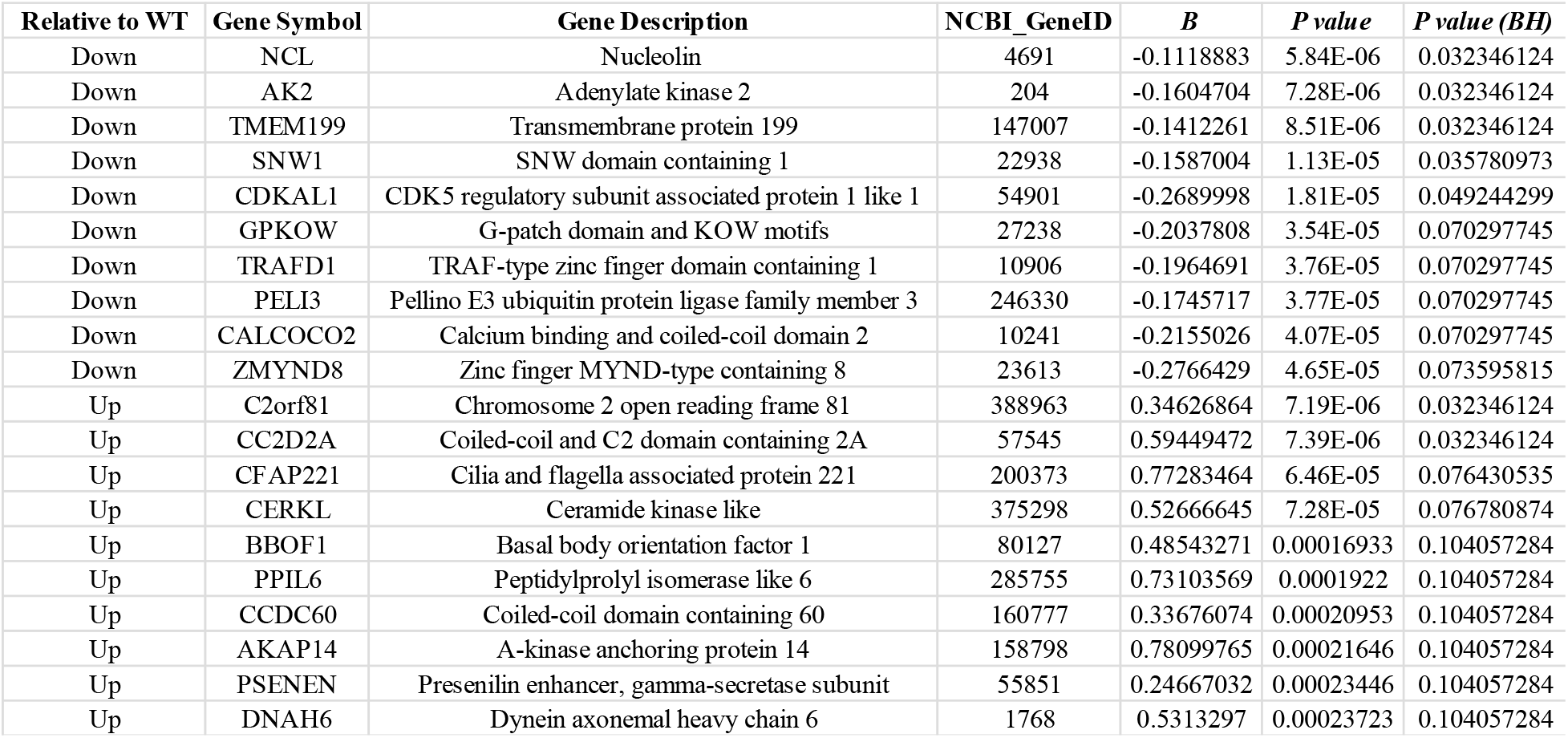
Wildtype vs CX4C-Mutant Differentially Expressed Genes

There were 354 biological processes enriched for gene transcripts that were decreased in the mutant CX4C virus compared to mock infection (Figure 2A, Supplemental Document 5). Biological processes enriched for decreased genes included response to cytokines, response to type I interferon, and RNA processing. Innate immune response biological processes including Nucleolin and many virus-signaling genes (e.g. BCL6, BATF and IRF2) were also decreased after mutant compared to wildtype infection.

**Figure 2.**
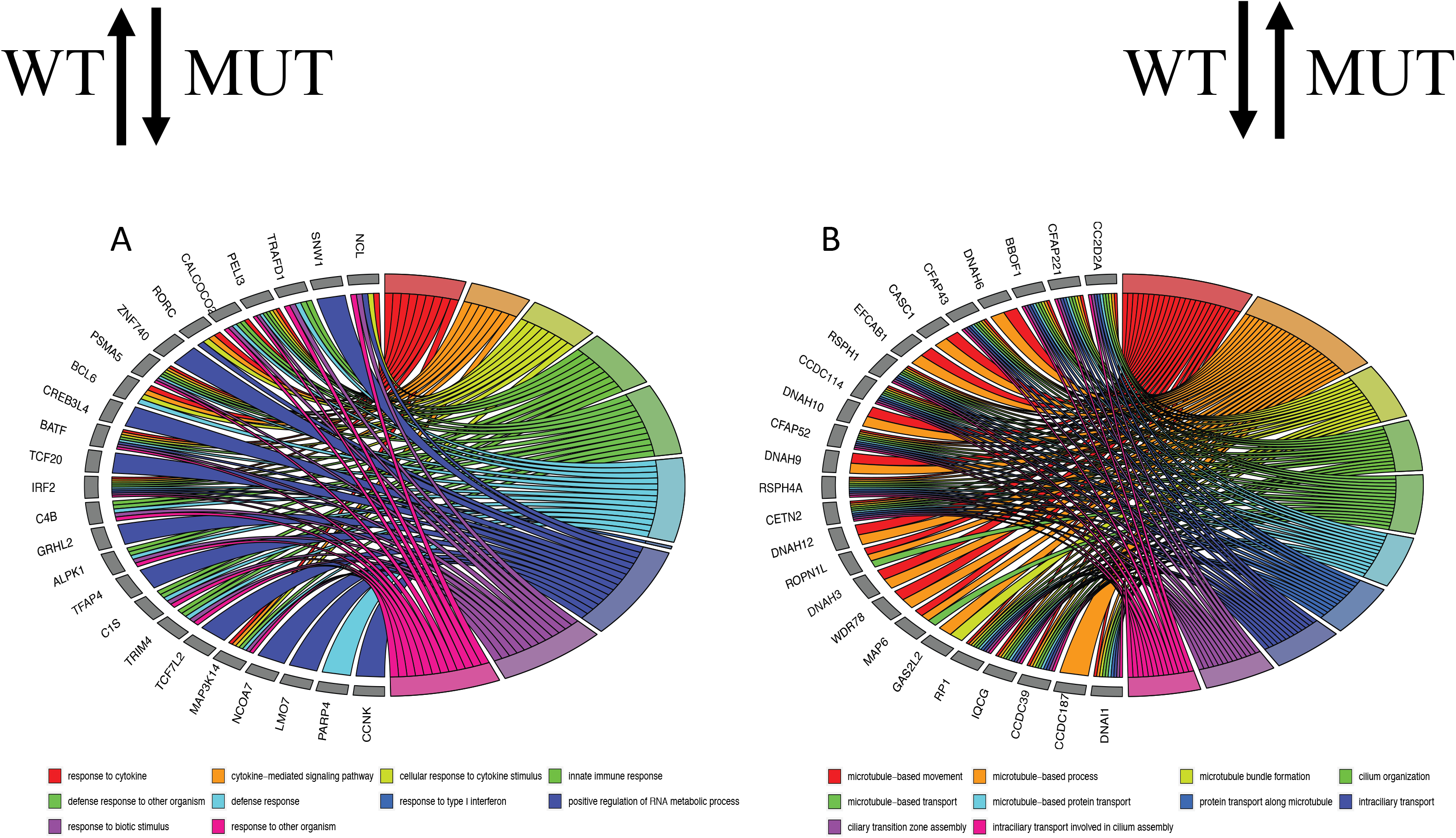
Changes in Biological Processes between wild-type and mutant RSV Infection. (A) Top ten biological processes, and associated genes, significantly enriched in transcripts increased after wild-type RSV infection compared to mutant (CX4C) infection. (B) Top ten biological processes, and associated genes, significantly enriched in transcript decreased after wild-type RSV infection compared to mutant (CX4C) infection. Colors represent the biological processes distinguished in the legend.

After multiple comparison correction, two gene transcripts were significantly increased after mutant CX4C RSV infection compared to wildtype virus (Table 2, Supplemental Document 4). The most significantly associated transcript, C2orf81, has no known function and the other transcript, CC2D2A, encodes a protein that forms a complex localized at the transition zone of primary cilia.

The 1067 transcripts increased (uncorrected p-value < 0.05) after mutant virus infection relative to wildtype infection resulted in 78 biological pathways that were significantly enriched (Figure 2B, Supplemental Doc 6). The biological processes were all associated with cilia and microtubule function and formation. These included transport, organization, and assembly. The genes associated with these pathways included many cilia-related genes (CC2D2A, CFAP221, CFAP43, CFAP52). Dynein-related genes (DNAH3, DNAH6, DNAH9, DNAH10, DNAH12) were also associated with enriched pathways. Taken together, mutating the CX3C domain resulted in a significant increase in cilia-related genes and biological processes.

### Transcriptional Changes with CX3CR1 Ligand

Recombinant G-protein soluble ligand representing the wildtype virus resulted in significant changes in transcript levels (Figure 3). Cilia-related genes (CC2D2A, CFAP221) were both significantly decreased after addition of G-protein ligand compared to mock treated cultures. Alternatively, Nucleolin was increased after treatment with G-protein ligand, although the relative fold-change was minor. Taken together, soluble G-protein effected transcript levels similarly to the differences seen between wildtype and CX4C mutant infections.

**Figure 3.**
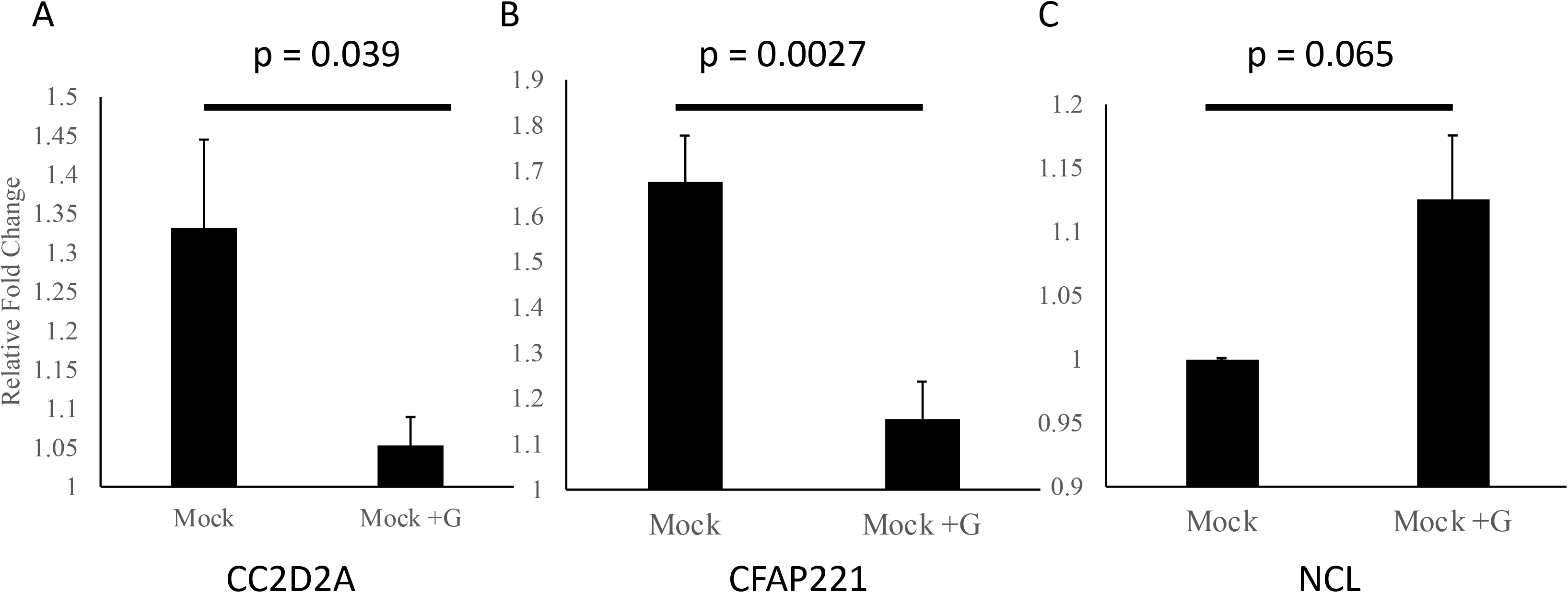
Effect of G-protein on Epithelial Cell Transcripts. Transcript levels of expression after treatment of epithelial cell cultures with recombinant G-protein for (A) CC2D2A, (B) CFAP331, (C) and Nucleolin.

### Ciliated Cells after RSV Infection

Immunohistochemistry of mock or RSV infected cells was performed in order to further understand the effect of RSV infection on ciliated cells (Figure 4A). Ciliated cells were found in both infected and non-infected cultures. Epithelial cells infected with wildtype RSV showed a significant decrease in the number of ciliated cells compared to mock infection (Figure 4B). Taken together, both mock and infected cells contained ciliated cells, but RSV infection decreased the percentage of ciliated cells relative to mock infection.

**Figure 4.**
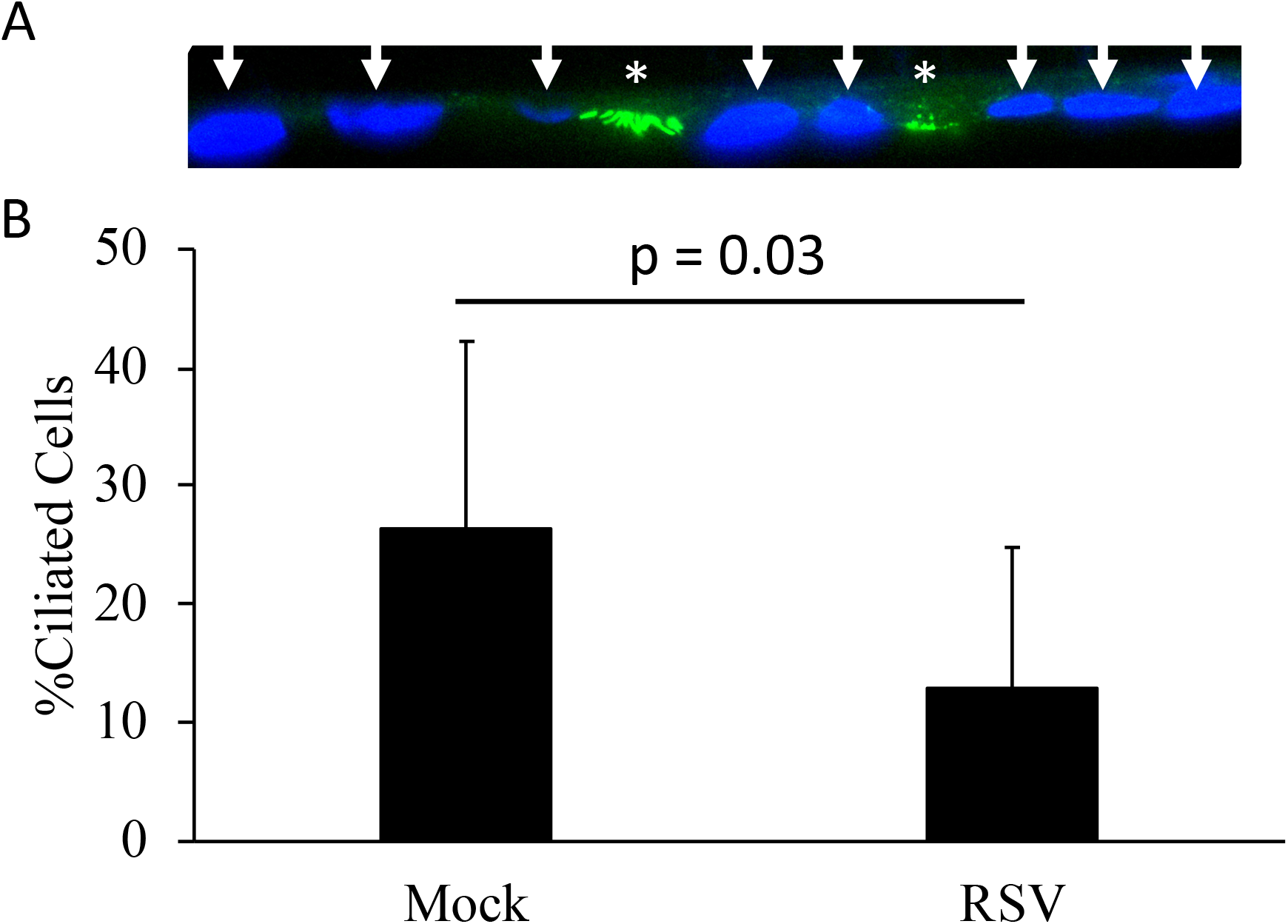
Ciliated Cells after RSV Infection. Quantification of the number of ciliated cells staining positive for acetylated tubulin per nucleus (dapi) in mock or RSV infected cultures.

## Discussion

Respiratory epithelial cells act as both the primary target for RSV and the primary defense against the virus. Therefore, it is unsurprising that so many genes and biological processes were altered during RSV infection. For successful replication the virus must first attach to the epithelium, enter the cell, dispense genetic material, take over host translation machinery, and use host nutrients create new genomes and protein, assemble, and release from the cell(1). During this process, the host epithelium recognizes the foreign object, produces anti-viral peptides, and signals to neighboring cells (including immune cells) of the infection(29). Therefore, the virus must both avoid host responses and hijack cellular processes in order to successfully reproduce.

Here we demonstrate that these underlying biological processes occurring during virus infection of adult primary epithelial cells grown at a physiological air-liquid interface. Given our experimental design, it is impossible to distinguish the transcriptional effects due to the host reaction to the virus and those caused directly by the virus. Both viral proteins and viral genetic material can interfere with biological processes by directly acting on host cellular machinery resulting in cellular signaling cascades that lead to changes in transcriptional regulation. It is of no surprise that these changes have to do with mechanistic ciliary function that removes foreign objects from the respiratory tract and the induction of Nucleolin, an RSV F-protein receptor(17), which would presumably provide a fitness advantage for the virus.

These introductory studies demonstrate a unique relationship between two host genes, CX3CR1 and Nucleolin, and two virus genes, G-protein and F-protein. These studies suggest that CX3CR1 engagement by the RSV G-protein CX3C motif results in intercellular signaling impacting Nucleolin expression. Although our experimental design did not allow us to discern any specifics in the signaling cascade, manipulating the cytokine pathway in order to provoke Nucleolin production would likely prove advantageous for the virus. Furthermore, these finding are consistent with the other viruses that manipulate host transcription in order to increase host proteins necessary to the virus(30).

Disruption of ciliary function has been described for many pathogens, although no morphology changes were seen after 24hrs post infection(31). A primary function of host immunity is the physical removal of foreign objects, including dead/infected cells, during an infection. It is important to note that the decrease in ciliated cells may be due to instability of the cilia and may have been disrupted during processing of the cultures. Regardless, it is clear that cilia, or the cells that express cilia, are affected by RSV infection. Given the role of cilia in the removal of foreign bodies, future studies will be needed to further characterize cilia function after RSV infection.

The CX4C mutant virus does not replicate at the level of the wildtype virus. Therefore, our statistical analysis required correction for variable levels of viral transcript. Although mathematical correction for viral transcripts levels using linear modeling have been continually shown to be accurate for these types of studies, it is not possible to completely distinguish Nucleolin induction or cilia dysregulation between wildtype and mutant based on CX3CR1 engagement from virus replication differences.

Furthermore, what effect disruption of the CX3CL1/CX3CR1 axis is having is unknown. It is entirely possible that the effect we see here is due to interference with autocellular signaling of CX3CL1 on CX3CR1. Crystal structures(32) suggest that engagement of CX3CR1 by RSV G-protein may be fundamentally different than fractalkine. Our studies cannot distinguish blocking of CX3CL1 binding from receptor activation. Moreover, our results do not rule out a general immune response to viral proteins. Future studies will be needed to establish the mechanism by which the G protein binding to CX3CR1 increases Nucleolin and decreases cilia-related transcripts.

It is important to note that these studies involved only a single strain of the A-subtype (A2). Given the large differences between subtype A and subtype B G-protein genes, and the fact that G protein has undergone significant genetic changes since the isolation of A2, future studies will need to determine if these transcriptional changes are consistent with RSV strains that have circulated recently and subtypes.

Taken together, RSV infection changes host gene expression and CX3CR1 appears to play a role in facilitating these changes. Our results suggest that RSV disrupts defense mechanisms while also increasing the expression of pro-viral proteins.

## Acknowledgements

We like to thank Jason Myers for his help in RNA sequences and data processing. We would also like to thank Dr. M.L. Moore for providing the viruses used in this study.

## Notes

This original work was supported by the NIH/NIAID HHSN272201200005C, the University of Rochester Pulmonary Training Fellowship NIH/NHLBI T32-HL066988, the University of Rochester HSCCI OP211341, the University of Rochester SAC Incubator Award, and the Department of Pediatrics, Emory University School of Medicine.

